# Data driven discovery and quantification of hyperspectral leaf reflectance phenotypes across a maize diversity panel

**DOI:** 10.1101/2023.12.15.571950

**Authors:** Michael C. Tross, Marcin W. Grzybowski, Talukder Z. Jubery, Ryleigh J. Grove, Aime V. Nishimwe, J Vladimir Torres-Rodriguez, Guangchao Sun, Baskar Ganapathysubramanian, Yufeng Ge, James C. Schnable

## Abstract

Hyperspectral reflectance data can be collected from large plant populations in a high-throughput manner in both controlled and field environments. The efficacy of using hyperspectral leaf reflectance as a proxy for traits that typically require significant labor and time to collect has been evaluated in a number of studies. Commonly, estimating plant traits using hyperspectral reflectance involves collecting substantial amounts of ground truth data from plant populations, which may not be feasible for many researchers. In this study, we explore the potential of data-driven approaches to analyze hyperspectral reflectance data with little to no ground truth phenotypic measurements. Evaluations were performed using data on the reflectance of 2,151 individual wavelengths of light from the leaves of maize plants harvested from 1,658 field plots of a replicated trial including representatives of 752 maize genotypes from the Wisconsin Diversity Panel. We reduced the dimensionality of this dataset using an autoencoder neural network and principal component analyses, producing 10 latent variables and principal components, respectively. A subset of these principal components and latent variables demonstrated significant repeatability, indicating that a substantial proportion of the total variance in these variables was explained by genetic factors. Moreover, correlations were observed between variables derived from the autoencoder network and principal components with molecular traits. Notably, the most relevant latent variable (LV8) showed a much stronger correlation with chlorophyll content (*R*^2^ = 0.59) compared to the most correlated principal component (PC2; *R*^2^ = 0.31). Furthermore, one latent variable exhibited modestly better performance than a partial least squares regression model in estimating leaf chlorophyll content (PLSR; *R*^2^ = 0.58, LV8; *R*^2^ = 0.59). A number of genetic markers in the maize genome were significantly correlated with variation in different latent variables in genome wide association studies. In a number of cases, significant signals in genome wide association studies were adjacent to genes with plausible links to traits expected to influence leaf hyperspectral reflectance patterns.

## Introduction

Mendel’s laws of genetics (law of segregation; law of independent assortment; law of dominance) were discovered through the analysis of qualitative traits (Weldon 1902; Biffen 1905). These laws created the field of genetics and have informed advances in plant breeding over the last century. Quantitative genetic approaches can use the combination of genetic marker data and trait measurements from large populations to identify individual genes or genomic segments associated with variation in target traits. However, collecting trait data from large plant field experiments is labor intensive and frequently represents the most expensive portion of quantitative genetics experiments of plant breeding efforts (Tibbs Cortes et al. 2021).

Advances in high throughput phenotyping technologies have the potential to reduce the cost of identifying genomic loci controlling traits of interest by either decreasing the resources required to score each plant/plot or reducing the marginal cost of collecting additional traits in parallel from a single field experiment (Fahlgren et al. 2015). Automated phenotyping strategies require solving two separate problems: collection of sensor data (e.g. RGB images, LIDAR point clouds, hyperspectral reflectance patterns, etc.) from plants of interest and using the collected sensor data to generate quantitative or qualitative estimates of specific traits (Yang et al. 2020; Furbank and Tester 2011)

Among many sensor modalities commonly used for high throughput plant phenotyping, spectrometers that collect leaf-level hyperspectral data in the visible, near-infrared, and shortwave-infrared regions (VIS-NIR-SWIR) are increasingly used to measure a wide range of plant chemical and physiological traits. Numerous studies have shown that VIS-NIR-SWIR can estimate leaf pigments, nitrogen content, water content, photosynthesis parameters, various metabolites, and nutrient contents (Yendrek et al. 2017; Silva-Perez et al. 2018; Vergara-Diaz et al. 2020; Chai et al. 2021). These spectrometers are portable, enabling data collection from field-grown plants, a clear advantage compared to many lab-only instruments.

Raw measurements of many individual spectral reflectance values are typically processed to generate predicted values for different traits of interest. Widely employed processing techniques for estimating known plant traits from raw spectral intensity values include narrow-band spectral indices, including the chlorophyll index (Wu et al. 2008) and the anthocyanin index (Steele et al. 2009), partial least squares regression (Burnett et al. 2021), and more recently approaches based on machine-learning and deep learning methodologies (Furbank et al. 2021). Both partial least squares regression and the set of machine learning methods described in (Furbank et al. 2021) are classified as “supervised” approaches, indicating that the models are trained using a population of data-points where both the spectral reflectance values and the true values for the trait of interest (labels) are already known.

Unsupervised methods can discern variation in sensor data gathered from various populations without the need to construct models targeting specific *a priori* traits or relying on labeled training data. This approach is particularly useful when dealing with datasets that exhibit high dimensionality, often with only a few degrees of variability. To boost predictive accuracy, implementing low-dimensional representations is advantageous as these highlight the fundamental characteristics of the data while filtering out unnecessary details. Techniques such as Principal Component Analysis (PCA) and Neural Networks (NNs) with auto-encoding are frequently utilized for this type of dimensionality reduction. While these methods might be abstract and sometimes challenging to interpret biologically, they are effective in uncovering hidden patterns in phenotypic and genetic variations (Ubbens et al. 2020).

PCA focuses on linear transformations to extract latent features, whereas NNs use a mix of linear and non-linear transformations. Studies by Wang *et al*. (2016) and Fournier and Aloise (2019) have empirically demonstrated the greater effectiveness of NNs over PCA in dimensionality reduction. In the realm of plant science, the efficiency of NNs has been confirmed by several researchers. Classifications and ordering of shape categories in strawberries using 2D images achieved heritabilites comparable to direct human measurement (Feldmann et al. 2020). Gage and coworkers demonstrated that quantitative latent phenotypes extracted from LIDAR point clouds in maize fields can exhibit heritabilities similar to hand measured traits (Gage et al. 2019).

Autoencoder neural networks are an unsupervised approach which reduce high dimensional data to a smaller set of latent variables that will, ideally, representing patterns of variation present in the original higher dimensional dataset (Wang et al. 2016; Rumelhart et al. 1985; Baldi 2012). Autoencoders comprise of two neural networks, an encoder and a decoder. The encoder takes as input, the values of all dimensions from a single sample in the dataset, reduces the dimensionality down to a configured amount of latent variables. The decoder then takes the latent variables as input and tries to reconstruct the sample to the original dimensionality. The reconstructed data from the decoder is then compared to the input of the encoder and the reconstruction loss is calculated. The loss is then backpropagated into the encoder and decoder networks to improve the parameters in the direction of better reconstructions. Through many iterations using numerous samples, the encoder is able to produce latent phenotypes that are representative of the original data.

Here we extract latent phenotypes from hyperspectral leaf reflectance sensor data in a maize diversity panel. We demonstrate how these latent phenotypes can be annotated and used as a proxy for traits with limited to no ground truth data.

## Materials and methods

### Field experiment and data collection

A previously described field experiment consisting of 1,680 plots subdivided into two complete replicates of a population of 752 maize inbred genotypes comprising a subset of the Wisconsin Diversity panel (Mazaheri et al. 2019), and a single repeated check genotype was planted on 6, May, 2020 at the Havelock Farm research facility at the University of Nebraska-Lincoln (40.852 N, 96.616 W) (Sun et al. 2022). Briefly, each plot consisted of two rows with approximately 20 plants per row. Rows were 7.5 feet long, with 30 - inch row spacing and 2.5 feet alley ways between sequential plots. Published plant-level phenotypes for the same experiment were taken from Mural *et al*. (2022). Hyperspectral reflectance data was collected on nine days over a 13 day period from July 8^*th*^ to July 20^*th*^, 2020. Hyperspectral reflectance was collected from a single fully expanded leaf from a representative plant per plot avoiding edge plants when possible using a benchtop spectroradiometer(FieldSpec4, Malvern Panalytical Ltd., Formerly Analytical Spectral Devices) with a contact probe, following a previously described protocol (Ge et al. 2019). A total of 2,151 reflectance values were collected per measurement measuring the proportion of light reflected in one nanometer increments from 350 to 2500 nanometers. Three spectral measurements were taken at the tip, middle and base of the adaxial side of each leaf. For each wavelength, values were averaged across the nine scans to generate a single final composite spectra for each plot sampled. A total of 1,665 plots were initially scanned. After averaging all scans for a given plot, plots with abnormal spectral were removed from the analysis, resulting in a final dataset of 1,658 plot level reflectance spectra.

Molecular leaf traits were collected from a subset of 243-318 of the same leaves from which hyperspectral leaf reflectance was also accumulated. Chlorophyll concentration (CHL), equivalent water thickness (EWT), leaf water content (LWC, %), and specific leaf area (SLA, *m*^2^/*kg*) was collected from these leaves using the methods adopted by Ge et al and Li et al. (Ge *et al*. 2019; Li et al. 2023). Briefly, CHL was measured using a handheld chlorophyll meter (MC-100, Apogee Instruments, Inc., Logan, UT), EWT was calculated using the formula: (*Freshweighto f leaves* − *Dryweighto f leaves*)/*lea f area*), LWC was calculated using; *dryweighto f leaves*/*Freshweighto f leavesX*100% and SLA was calculated using; *lea f area*/*dryweighto f leaves*. Phosphorus (P), nitrogen (N), potassium (K), magnesium (Mg), calcium (Ca), sulphur (S), iron (Fe), manganese (Mn), boron (B), copper (Cu) and Zinc (Zn) were quantified from dried leaf samples by a commercial provider (Midwest Laboratories, Inc., Omaha, NE).

A comparable set of hyperspectral reflectance and molecular leaf trait data was collected for a sorghum population grown in field conditions and under different nitrogen treatments as previously collected and described in Grzybowski *et al*. (2022). A separate experiment using genotypes from the same sorghum association panel grown in a controlled greenhouse environment was described in Tross *et al*. (2021), and the data collected was previously described in Ge *et al*. (2019).

### Dimensional Reduction

Dimensionality of the 1,658 plot-level hyperspectral reflectance values was reduced using both principal component analysis (PCA) and a trained autoencoder neural network. For PCA, the values were reduced to 10 principal components using the scikit-learn package (Pedregosa et al. 2011). These 10 components were sufficient to summarize 99% of variance in the dataset. An autoencoder architecture was implemented in keras (v2.8.0)(Bank et al. 2020; Chollet et al. 2015). The empirically determined network architecture consisted of an encoder with five dense layers with 2,151, 2,200, 3,000, 2,024 and 10 neurons respectively and a decoder with five dense layers with 1,024, 1,536, 2,500, 2,500 and 2,151 neurons (Figure S1). A scaled exponential linear unit (SELU) activation function was employed for all dense layers, with the exception of the final dense layer of the decoder network which employed a tanh activation function. Both the encoder and decoder were trained using a mean absolute error loss function, the standard gradient descent optimizer, and a learning rate of 0.1. The raw set of 1,658 plot-level hyperspectral reflectance values was split 5:1 into training and validation data. Autoencoders were trained for up to 1,000 epochs, or until 100 epochs passed without further improvement, whichever came first. The final autoencoder described in this manuscript was trained for 413 epochs before stopping based on a lack of further improvement.

To compare autoencoder latent variables to current state of the art methods for chlorophyll estimation, a scikit-learn implementation of a partial least squares regression model (Helland 1990) was trained using ground truth chlorophyll content measurements from 318 plots. The model was evaluated using five fold cross validation. The importance of hand-measured traits in predicting latent variables was derived using the built in feature importance function of the random forest model (Breiman 2001; Ho 1995) implemented in scikit-learn. Feature importance is referred to as the mean decrease in impurity (MDI)(Louppe et al. 2013). MDI calculates how much each feature contributes to organizing the data into more homogeneous or pure groups at each split in the decision tree nodes.

### Quantitative Genetic Analyses

Repeatability of plant phenotypes in this study was calculated using the equation 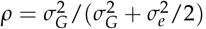, where 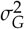 is the total amount of variance explained by genetics, and 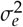 is the total amount of residual variance. Variance components were derived fitting a linear model with the formula *y*_*i*_ = *μ* + *t*_*i*_ + *e*_*i*_, where *y*_*i*_ is the mean value of the genotype, *μ* is the overall mean, *t*_*i*_ is the effect of genotype i, and *e*_*i*_ is the residual error of genotype i. Linear models were fit to each dataset using software package lme4 (v1.1-23) (Bates *et al*. 2015).

Genome wide association studies were conducted using the mixed linear model approach implemented within the GEMMA software package (v0.98.1) (Zhou and Stephens 2012) with a set of 16.6 million segregating genetic markers, a subset from the set of genetic markers published in Grzybowski et al. (2023), (Grzybowski et al. 2023) and filtered to include only those with a minor allele frequency *geq* ≤ 0.05 and a proportion of heterozygous calls ≤ 0.05. A total of three principal components of variation in the genetic marker data were calculated using PLINK (v1.90b4) software package (Purcell et al. 2007), and incorporated as covariates into the model employed for genome wide associations. The threshold for statistical significance in this study – 2.20 x 10-8 – was determined by applying the Bonferroni correction to the estimated 2,269,711 effective number of independent statistical tests represented by the 16.6 million markers employed in this study. The effective number of independent markers was calculated by pruning those initial 16.6 million markers using a sliding window of 500 bp, step size of 100 and removing all SNPs above a linkage disequilibrium of 0.2 using PLINK.

An eQTL(expression Quantitative Trait Loci) mapping analysis as described in Torres-Rodriguez *et al*. (2023) was conducted using the rMVP (V1.0.6)(Yin et al. 2021) implementation of the mixed linear model. Briefly, gene expression for the Zm00001eb29707 gene model across genotypes was transformed using the Box-Cox method (Osborne 2010) and the genetic marker dataset was processed as previously described. A kinship matrix generated using the VanRaden method (VanRaden 2008) and three principal components of variation in the genetic marker dataset was used as covariates in the analysis.

## Results

### Hyperspectral leaf reflectance values

The proportion of light hitting the adaxial surfaces of maize leaves harvested from a replicated field study of more than 700 maize genotypes which was reflected varied between 0.2% and 52% for individual one nanometer wide bands of light between 350nm and 2500nm (3,566,358 observations, 2,151 individual wavelengths * 1,658 leaves). Pairwise correlations between the intensity with which different one nanometer wide wavelengths of light were reflected by different maize leaves in the dataset ranged from -0.1 to 0.99 (Spearman’s rho) with blocks of wavelengths exhibiting high correlations (Figure 1A). The proportion of the variance in the reflectance of individual one nanometer wide wavelengths, which could be explained by differences between maize genotypes ranged from 0 to 0.45 (Figure 1B).

**Figure 1.**
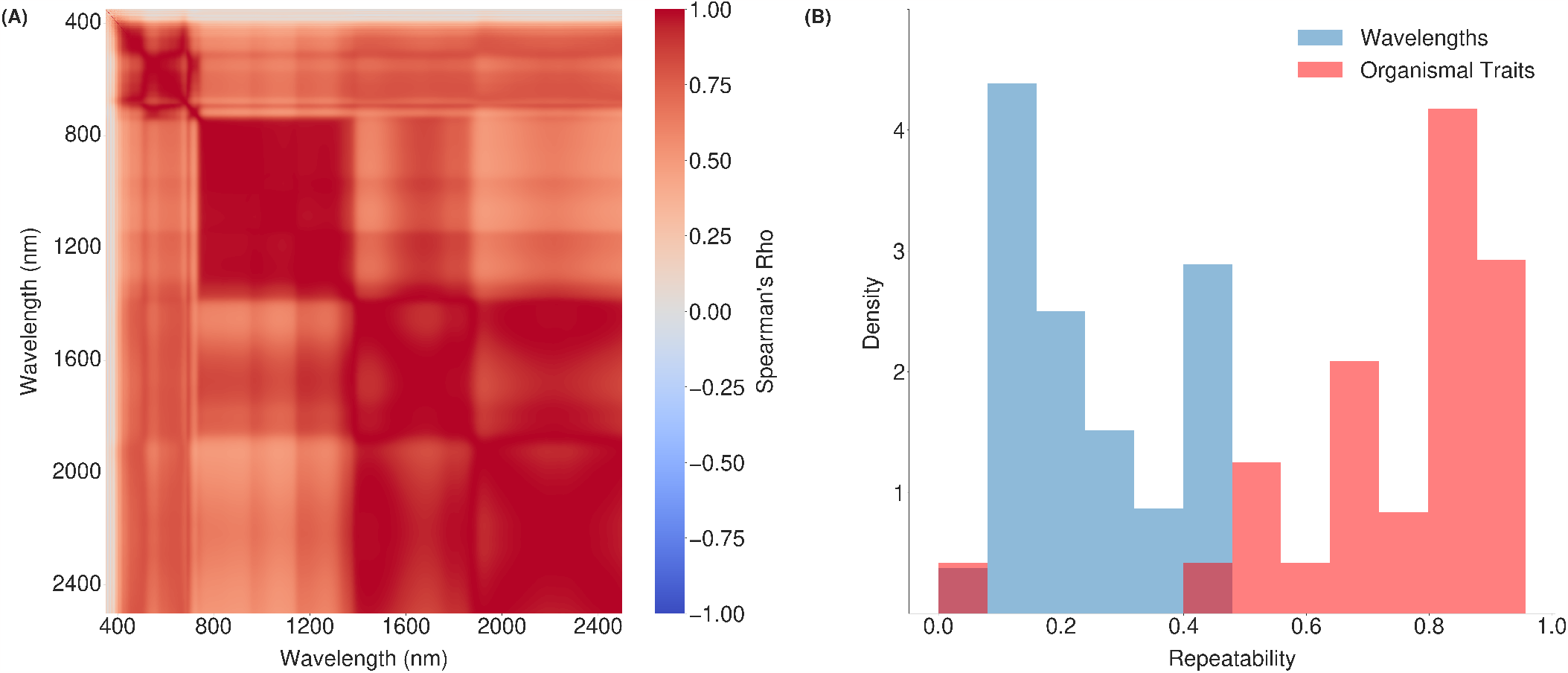
Correlations among the reflectance values of individual wavelengths and the proportion of variance attributable to differences between maize genotypes. (A) Spearman’s rho values across 1,658 maize leaves measured in a 2020 field experiment for all possible pairwise combinations of the proportion of light reflected for individual one nanometer wide wavelengths between 350 nm and 2,500 nm. (B). Comparison of the distribution of repeatability values – defined here as the proportion of variance attributed to differences between replicates of the same genotype in a single environment – for the reflectance of 2,151 individual one nanometer wide wavelengths and for a set of 30 hand measured traits scored from maize plants in the same field experiment Mural *et al*. (2022).

The substantial correlations observed among the reflectance values of many individual wavelengths of light suggested the potential to summarize leaf reflectance using a smaller number of variables. Variation in reflectance across all 2,151 individual wavelengths were summarized using autoencoders trained to summarize individual leaf reflectance spectra between one and 20 latent variables, and then reconstruct the original 2,151 variable data from the smaller number of variables. The architecture of these autoencoders differed only in the number of variables passed from the encoder to the decoder (see Methods). The resulting models were assessed in two ways: first, by the reconstruction loss observed on validation data not used to train the autoencoders, and second, by the correlation of the autoencoder variables with a set of 15 molecular leaf traits we quantified from maize plants in the same field experiment. The five lowest minimum reconstruction losses on the validation data of 20 trained models were 0.0122, 0.0132, 0.0128, 0.0129 and 0.0125 for models having five, seven, 10, 11 and 17 latent variables respectively (Figure S2). The maximum of maximum correlations across each latent variable with molecular traits was derived for models having five, seven, 11 and 17 latent variables. The correlation values were 0.51, 0.59, 0.66, and 0.64 respectively. Based on a combination of minimizing reconstruction loss and maximizing correlation with molecular traits measured from the same field experiment, the 10 latent variables model was selected for the analyses presented below. The final trained encoder was used to summarize leaf reflectance data from each of 1,658 plots as 10 total latent variables. The repeatability of four of these 10 latent variables exceeded that of any individually measured wavelength. The highest observed repeatability of a latent variable was 0.64, while the highest observed repeatability of an individual wavelength was 0.45 (Figure 2). A control which employed principal component analysis to summarize the same dataset produced 10 PCs among the first 10 with repeatability 0.5 (two less than the autoencoder approach) and a maximum repeatability of 0.59 (0.64 for the autoencoder model (Figure S3). In several cases, latent variables with low repeatability appeared to represent variation between leaves analyzed on different days (Figure S4). The latent variable most correlated with chlorophyll content exhibited a correlation of (*R*^2^ = 0.59)(Figure 2F), which was which was much greater than the highest correlation observed between any of the first 10 principal components and chlorophyll (*R*^2^ = 0.31) (Figure S6). It matched, and in fact modestly exceeded, the predictive accuracy of supervised models (partial least squares) individually trained on different 80% subsets of the same 318 ground truth chlorophyll measurements used to evaluate both models (*R*^2^ = 0.58) (Figure 2E).

**Figure 2.**
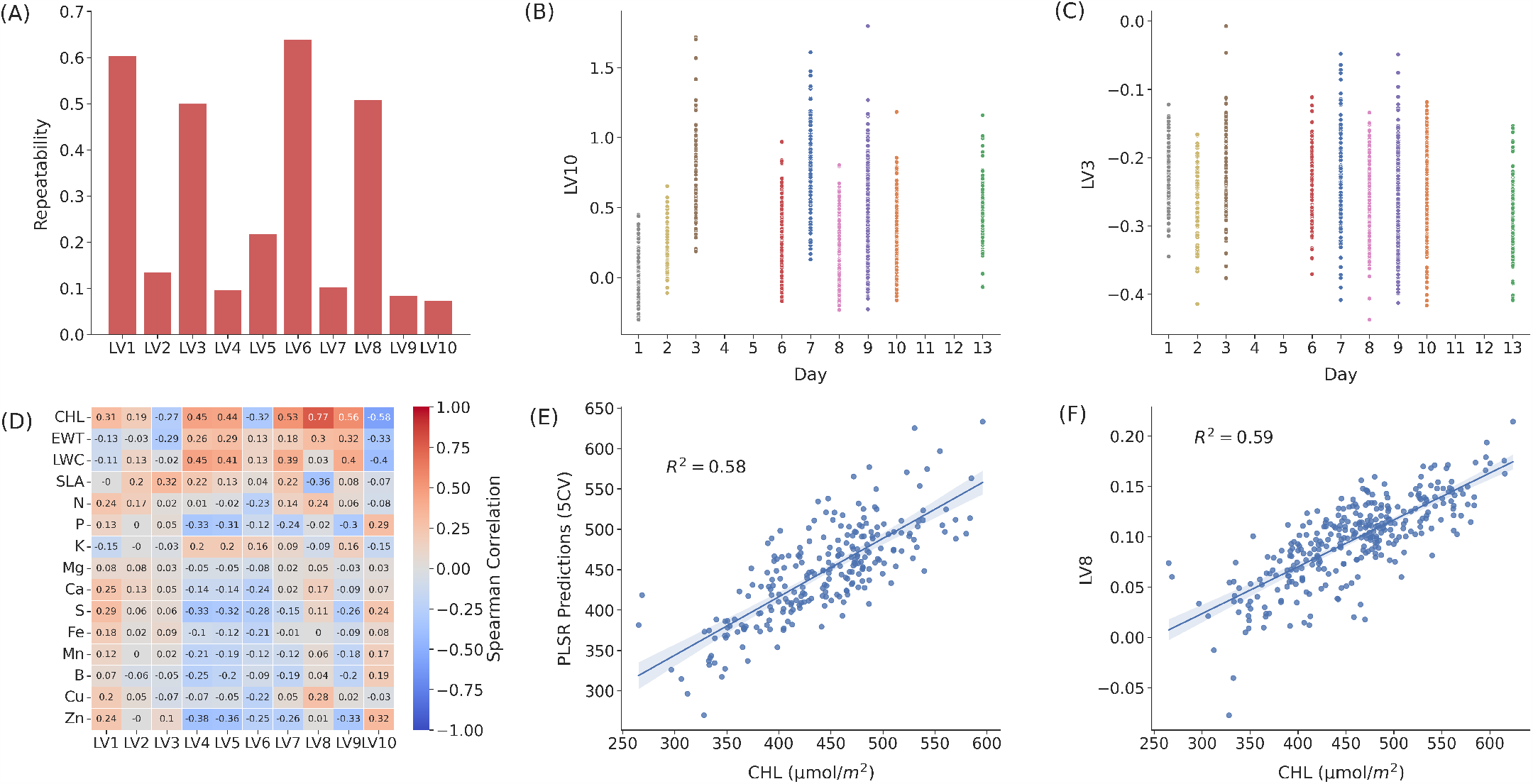
Association of latent variables with genetic, environmental, and plant phenotype factors. (A). Comparison of the repeatability of each of the 10 latent variables derived from hyperspectral leaf reflectance measured in this study. (B). Latent variable 10 value of the leaf reflectance compared with the day of collection of leaf reflectance for each plot. (C). Latent variable three value of the leaf reflectance compared with the day of collection of the leaf reflectance for each plot. (D). Associations measured by Spearman’s rho between individual latent variables and ground truth measurements for 15 traits each scored in subsets of between 243 and 318 maize plots from which leaf reflectance data was also collected in 2020. (E). Association between observed chlorophyll content and five fold cross validation predictions from partial least squares regression model for all experimental plots. (F). Association between observed chlorophyll content and latent variable eight of the leaf reflectance values for all experimental plots.

The transferability of the autoencoder derived variables with plant traits was assessed using two sorghum leaf reflectance datasets and associated ground truth data (Grzybowski et al. 2022; Wijewardane et al. 2023). The pre-trained encoder described above was used to summarize variation from hyperspectral leaf reflectance collected from 321 sorghum plants grown under greenhouse conditions in 2018, 132 sorghum plants grown in the field in 2020 under sufficient nitrogen conditions, and 136 sorghum plants grown in the field under nitrogen deficient conditions in 2020. The same latent variable continued to exhibit correlations with ground truth chlorophyll measurements in greenhouse grown sorghum (*R*^2^ = 0.24), field grown sorghum with optimal nitrogen conditions (*R*^2^ = 0.61), and field grown sorghum under nitrogen deficient conditions (*R*^2^ = 0.64) (Figure 3).

**Figure 3.**
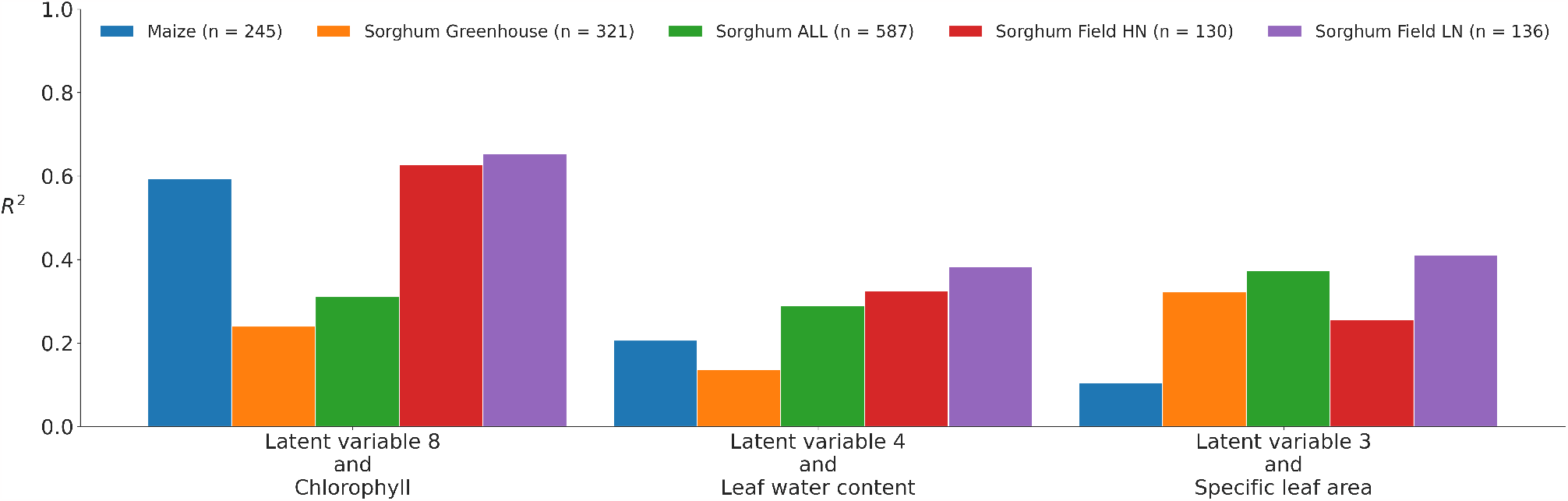
Associations between latent variables of leaf hyperspectral reflectance of sorghum populations with the same traits across species and years. All latent variables are derived from the same autoencoder model trained on hypersepctral reflectance data from a maize species. Latent variables derived from maize field reflectance data (2020), sorghum leaf reflectance field data (2020) (Grzybowski et al. 2022) and greenhouse data (2018) (Wijewardane et al. 2023), predicted from a model trained on leaf reflectance in maize (2020) were all associated with the same molecular traits.

### Latent variables capture information on variation in organismal traits

Random forest models were trained to predict latent variables from a suite of 30 traits measured from maize plants in the same field experiment to determine if latent variables were associated with variation in other traits of interest in the same maize population. Feature importance, quantified as the mean decrease in Gini impurity, was assessed for each organismal trait in random forest models trained to predict each latent variable. This produced an assessment of which plant traits contained significant information about the value of each latent variable summarizing variation in leaf light reflectance. Leaf width exhibited the highest five fold mean decrease in impurity (0.12) for latent variable three, with the second largest decrease exhibited by the trait number of branches per tassel (0.08) (Figure 4). Severity of southern rust legions, a leaf pathogen observed later in the growing season in the same field leaf reflectance data was collected from, was the most important organismal trait in predicting latent variable eight (Figure S7). Flowering time (number of days to pollen and number of days to silking) were more important predictors of the two latent variables with the highest repeatabilities (Figure 2A), and were latent variables one and six (Figure S7).

**Figure 4.**
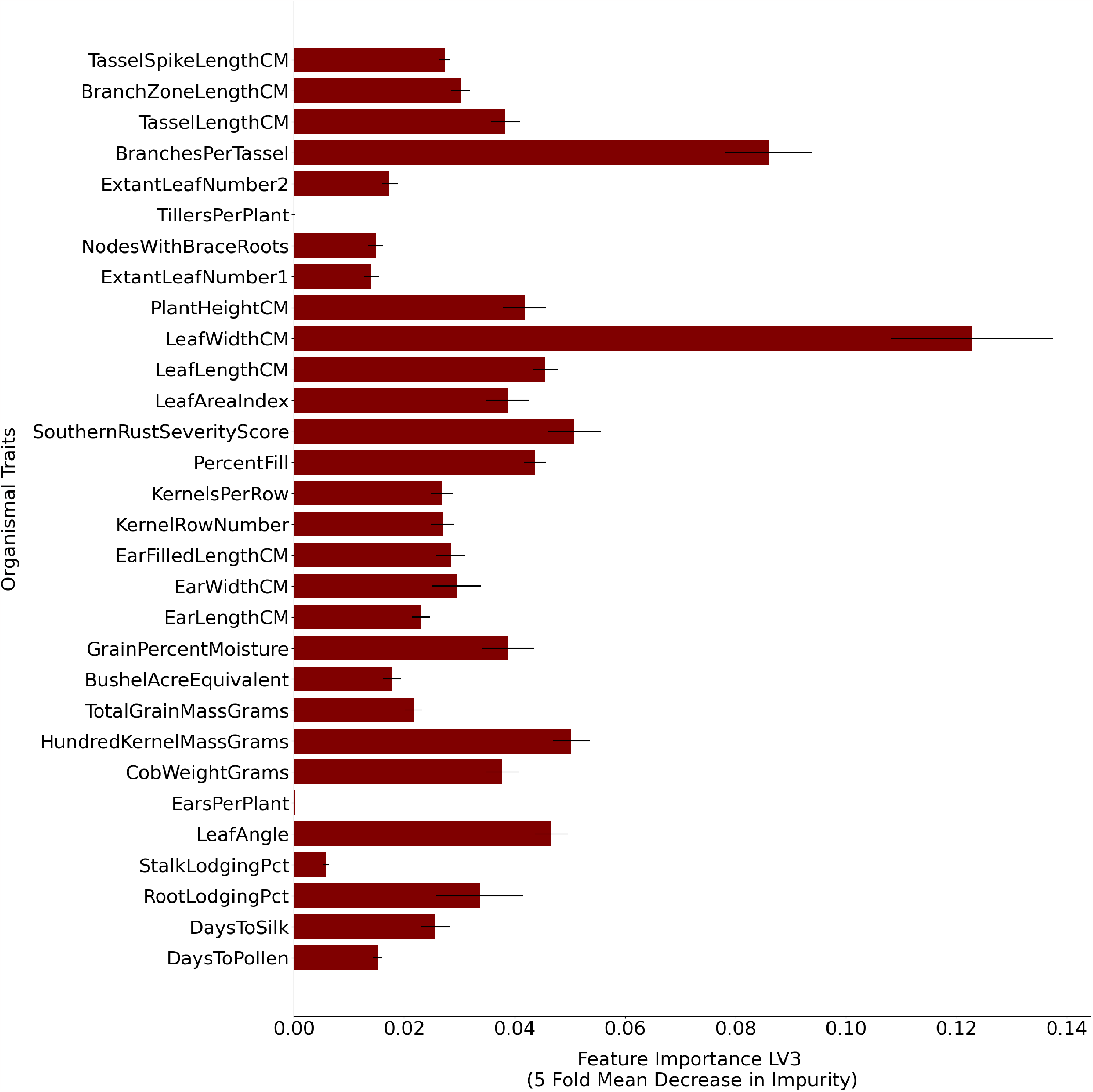
The relative importance of 30 hand measured traits in predicting latent variable three of the hyperspectral leaf reflectance values. This importance is derived from the mean decrease in impurity of each node in the decision trees within a trained random forest model, attributed to each feature (trait). X axis indicates the five fold mean decrease in impurity calculated for each trait. Error bars represent standard error across the five folds for each trait. Y-axis indicates the 30 hand measured traits used to predict the latent variable three.

### Linking latent variables to causal genes via genome wide association

Many latent variables were not strongly associated with molecular or organismal traits. Genome wide association studies were conducted on all latent variables to better understand what types of mechanisms might underlie the variation captured by each variable. Most latent variables exhibited statistically significant association with at least one genetic marker in the maize genome (Figure S8). Latent variable three had a significant marker that was located 15,811 base pairs downstream from Zm00001eb134990. The Arabidopsis ortholog of this gene, CYCD5;1 (AT4G37630) is believed to play a role in controlling edoreduplication during leaf development, a process associated with trichomes and other specialized protruding cells from the leaf surface (Sterken et al. 2012). A genetic marker 7,713 base pairs upstream of Zm00001eb297070 was significantly associated with variation in latent variable five (Figure 5G). Notably this signal was also only 988 base pairs away from the peak SNP of an eQTL associated variation in the expression of that same gene (Zm00001eb297070) (Torres-Rodriguez et al. 2023) in mature leaf tissue (Figure 5H). A significant hit for latent variable six was located within the annotated gene model for Zm00001eb434330 (Figure 5F), a gene expressed primarily in developing leaves in maize (Stelpflug et al. 2016; Hoopes et al. 2019).

**Figure 5.**
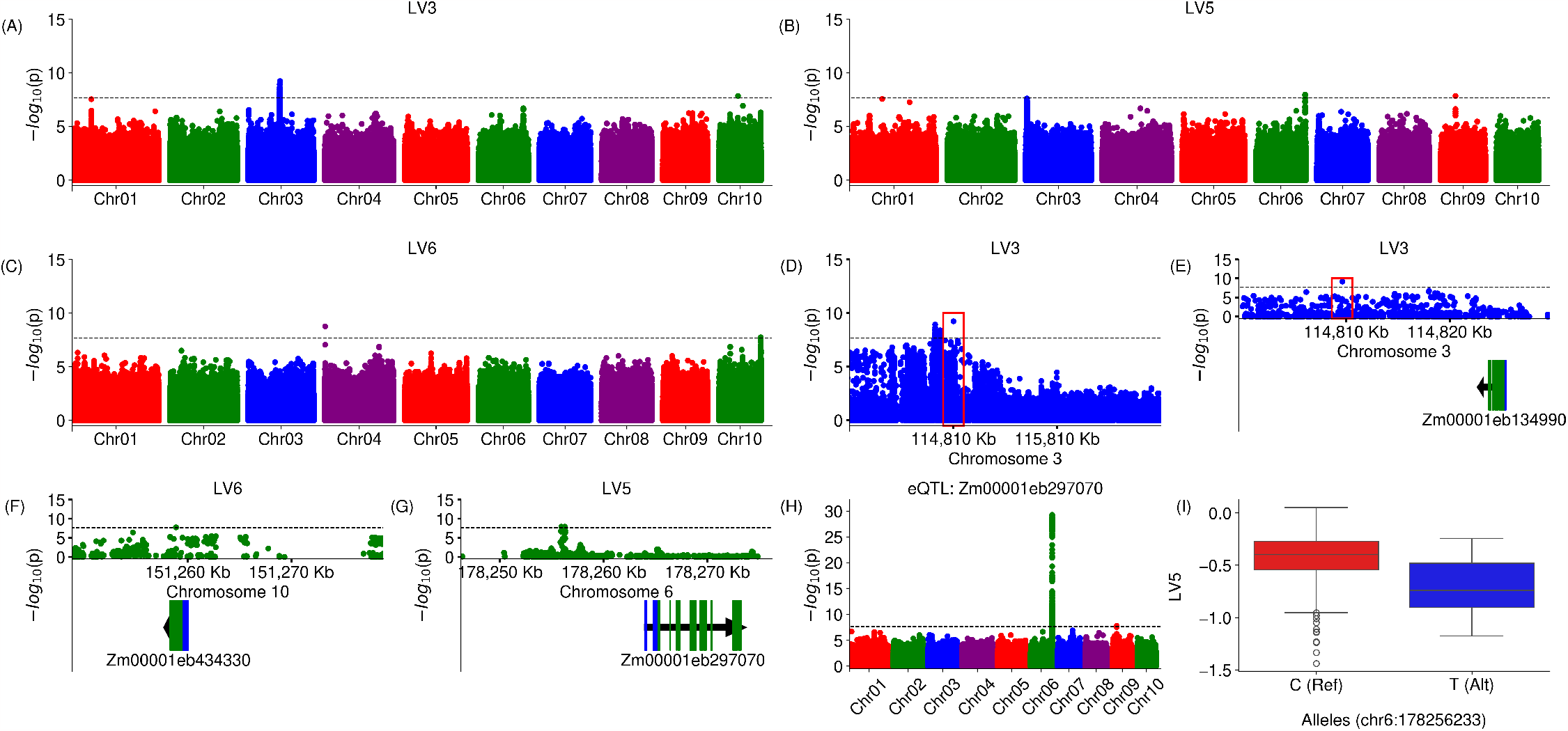
Genetic markers significantly associated with latent variables three, five, and six of autoencoder trained on reflectance data. Each point indicates the statistical significance of a marker (y axis) and its exact position on the genome (x axis). The dashed black lines indicate the statistical threshold cut off –2.20 x 10-8 – which was derived from an alpha value of 0.05 with a Bonferonni adjustment for 2,269,711 effective genetic markers. Annotated black triangles indicate the position of the nearest genes and gene models. Genome wide association studies of: (A) Latent variable three; (B) Latent variable five; (C) Latent variable six; (D) Zoomed in region of the significant markers on chromosome three of latent variable three starting at **113**,**809 kb** to **116**,**809 kb**. Red bounding box indicates the region of the most significant SNP; (E) Zoomed in region of the the significant markers on chromosome three of latent variable three and the nearest genes and gene models starting at **114**,**799 kb** to **114**,**829 kb**. Red bounding box indicates the region of the most significant SNP; (F). Zoomed in region of the of the significant markers on chromosome 10 of latent variable six and the nearest genes and gene models. (G). Zoomed in region of the significant markers on chromosome six for latent variable five and the nearest gene models; (H) eQTL analysis of the Zm00001eb297070 gene model; and (I) Comparisons of the distributions of latent variable 5 for genotypes that are homozygous for the ‘C’ reference allele versus the ‘T’ alternate allele.

## Discussion

The cost and throughput of collecting accurate measurements of plant traits across large field experiments is increasingly the rate limiting step in both plant quantitative genetics research and plant breeding. Approaches that substitute sensor data and prediction models for direct human measurements of traits have been adopted for some applications and show promise in others. However, common approaches to training models to predict traits for sensor data using supervised models require large and expensive datasets to train, making them inaccessible to many researchers working on specialty crops, genetic models, or previously poorly studied traits. Here, we aimed to quantify traits using a high dimensional hyperspectral leaf reflectance dataset combined with data-driven approaches which have the potential to mitigate some of the logistical challenges of supervised training models.

A greater proportion of latent variables produced by autoencoder based summaries of leaf reflectance data exhibited repeatabilites >0.4 than did principal components calculated from the same leaf reflectance dataset (Figure S3). However, repeatable traits can still be of limited utility for plant breeding and genetics if those traits are not linked to known plant properties. One autoencoder derived latent variable captured variation in chlorophyll content with an accuracy which matched or modestly exceeded that of a supervised model trained with labeled data while none of the top ten PC approached the performance of the supervised model (Figure 2 & Figure S6). For many large quantitative genetics studies, variation in phenotypic measurements due to temporal variation in the time different plants or plots are scored creates additional non-genetic variance and reduces power to either identify causal genes or build trait prediction models. We observed that several latent variables appear to reflect day of collection and were less associated with differences between genotypes (e.g. repeatability), while other latent variables were primarily associated with genetic effects (Figure 2B & C, Figure S4). This variance partition-like property may explain, at least in part, the greater correlation of some latent variables with other, ground truth, plant phenotypes.

Beyond chlorophyll, a trait which can already be predicted with high accuracy with a number of linear models trained on large datasets, the strongest correlation of any of the latent variables calculated in this study with a panel of molecular traits was approximately R^2^=0.2. Similarly, low correlations were observed with a panel of whole-plant phenotypes (Figure S9). However, linear regression may not capture non-linear relationships between traits, or cases where a single latent variable reflects variation across multiple molecular or whole plant traits. Feature importance values calculated from random forest models, which can capture both non-linear relationships and the influence of multiple traits single latent variables, enabled the identification of whole plant phenotypes, including leaf width, flowering time, and susceptibility to a specific foliar pathogen, associated with multiple individual latent variables (Figure S7). However, it must be noted that this approach was unsuccessful for a number of latent variables. Success depended on access to large datasets of conventionally scored traits from the same populations from which leaf reflectance data was collected, potentially reducing the logistical advantages of this approach relative to training supervised models. Another potential strategy for linking autoencoder derived latent variables to known plant properties is via quantitative genetics (Ubbens et al. 2020). If a given latent variable is associated with a variable in multiple genes known to control a specific plant trait of interest, this would serve as significant evidence that the latent variable reflects variation in the same trait. We were successful in identifying one or more numbers of genomic intervals which were significantly associated with variation in seven out of 10 latent variables. In at least one case, a genome wide association study hit was associated with genes with plausible links to leaf-reflectance related phenotypes. A genetic marker significantly associated with latent variable three (Figure 5A) was identified 16 kilobases from a maize gene whose Arabidopsis ortholog is associated with leaf development and differential organ growth in different environments (Sterken et al. 2012). In another case a genetic marker associated with latent variable five (Figures 5B & 5G) was also associated with cis-eQTL for a variable in the expression of an adjacent (approximately one kilobase distant) gene (Figure 5H). However, this approach was limited by both the number of GWAS hits identified per latent variable, and the relatively modest number of maize genes linked with high confidence to roles in determining plant phenotypes. The former issue can potentially be addressed in the future by collecting hyperspectral leaf reflectance data from larger populations, experiments with higher levels of biological replication within a single environment, and/or across greater numbers of environments. Each of these would increase our power to identify significant associations in genome wide association studies. The capacity of encoders trained on a single environment to continue to accurately reflect variation in the same plant phenotype across datasets collected in multiple environments and from multiple species (e.g. maize and sorghum) suggests that this approach may indeed be feasible (Figure 3). Employing autoencoders for dimensionality reduction requires a substantially greater amount of user time and input than principal component analysis. Data conversion, network architecture design, hyperparameter tuning, and access to the necessary types and scale of computer resources are all barriers of entry relative to current widely used methods of dimensionality reduction in plant biology applications, including principal component analysis, and current widely used supervised classification models such as partial least squares regression. In addition, while the collection of leaf hyperspectral reflectance data for large plant populations is less labor intensive than manual scoring of large panels of plant traits from the same population, the costs of the necessary equipment are high and the labor requirements are nontrivial. However, current rapid advances in robotics and imaging technologies have the potential to address the second challenge, while improved artificial intelligence and machine learning frameworks may address the first. If so, the approaches described here may provide significant utility in assisting plant geneticists and plant breeding in extracting the maximum amount of useful information from these new data types.

## Code and Data availability

The code used in this study is available at https://github.com/mtross2/autoencoder_hyperspec_ref The hyperspectral dataset, molecular leaf traits, and weights for autoencoder model generated in this study is available at https://doi.org/10.6084/m9.figshare.24808491.v1

## Acknowledgments

The authors thank Christine Smith and Mackenzie Zwiener for supervising and designing the 2020 maize and sorghum fields described in this manuscript. We thank Nuwan K. Wijewardane and Abbas Atefi for both expert guidance and help in the collection of the hyperspectral reflectance data described in this study. We thank Nikee Shrestha for assistance in curating datasets. We thank Addie Thompson and Linsey Newton for packaging and shipping the seeds during the 2020 coronavirus lockdown that enabled the maize field experiment described in this study. We thank Nathaniel Pester, Leighton Wheeler, Sierra Conway, Isaac Stevens, and Luke Micek for assisting in field maintenance and data collection.

## Author Contributions

MCT and JCS conceived of the project. MWG, GS, AVN, YG generated, assembled, and quality controlled data. MWG, TZJ, BG and JCS designed and advised on analysis methods. MCT, RJG, AVN, JVTR conducted analyses. MCT, JCS wrote the first draft of the manuscript. RJG, JVTR, TZJ, BG, YG contributed significant additional content during the revision of the manuscript. All authors read and approved the final version of the manuscript.

## Supplementary materials

**Figure S1.**
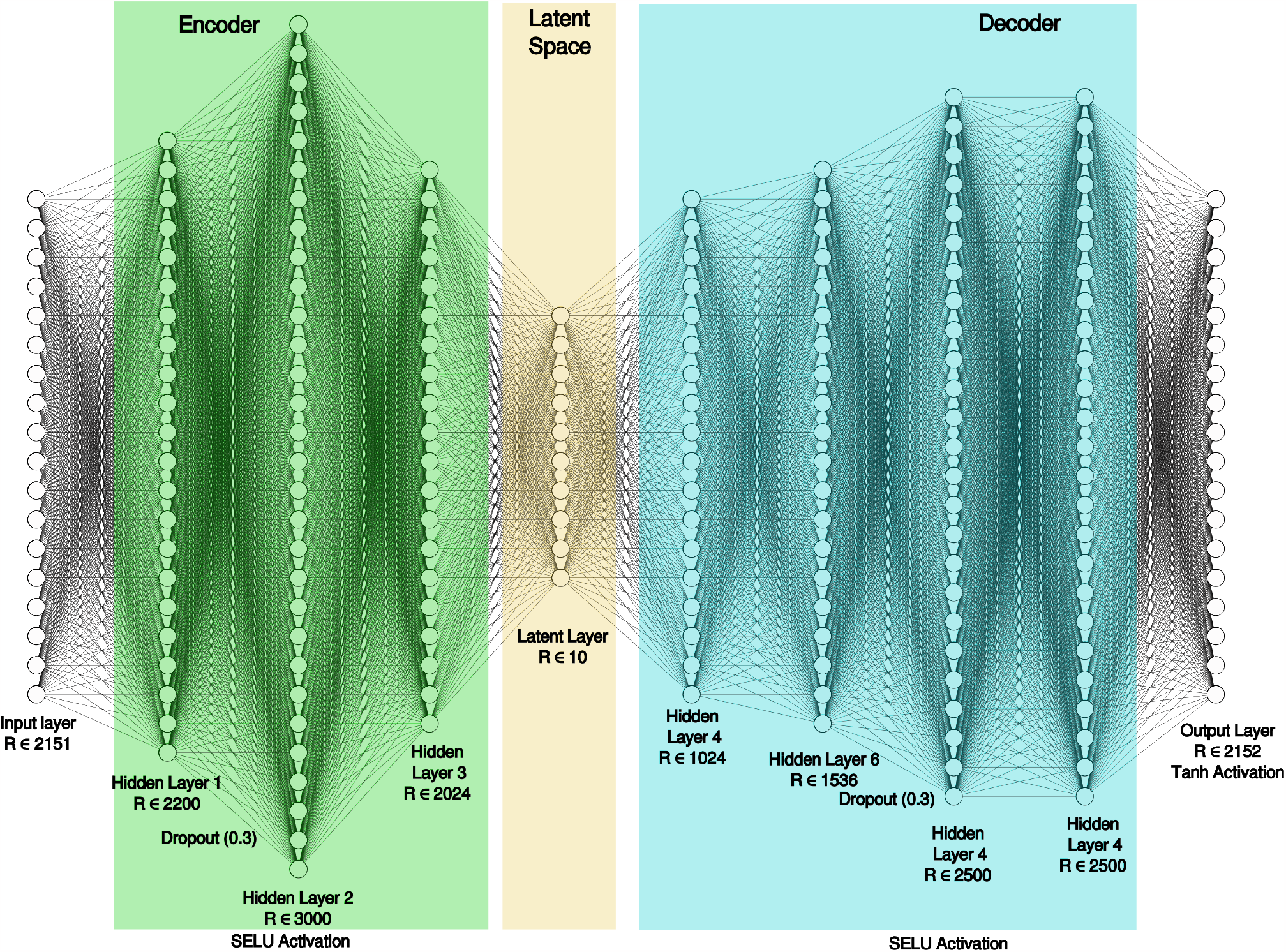
Architecture and hyper-parameters of auto-encoder neural network employed in this study. The encoder consisted of five dense layers with 2,151, 2,200, 3,000, 2,024, and 10 neurons respectively, and a decoder with five dense layers with 1,024, 1,536, 2,500, 2,500 and 2,151 neurons. Each dense layer employed a scaled exponential linear unit (SELU) activation function, except for the final dense layer which employed a tanh activation function.

**Figure S2.**
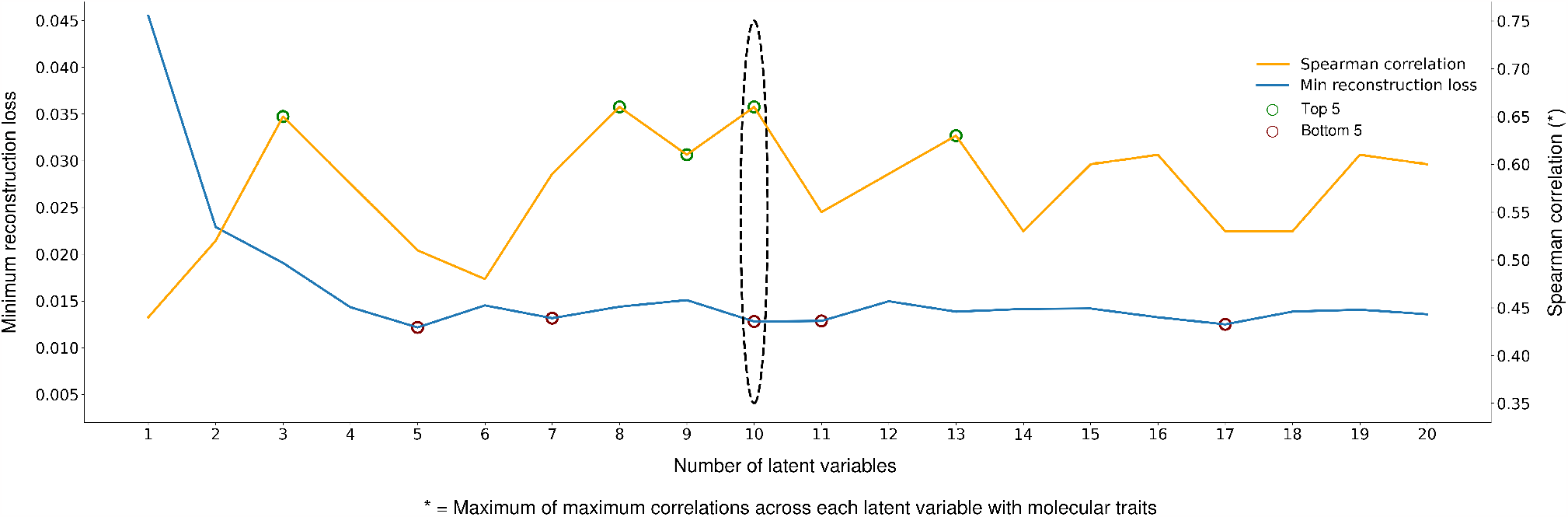
Selection of the number of latent variables used in auto-encoder architecture. Blue solid line indicates the minimum reconstruction loss with the incremental increases in number of latent variables, ranging from 1 to 12. Red circles indicate the bottom 5 minimum reconstruction losses for all models. Yellow line indicates the maximum of maximum correlations across each latent variable with molecular traits with incrementing number of latent variables. Green circles indicate the top 5 Spearman correlation values across all models. Dotted black oval indicates the optimum number of latent variables based on having a top 5 Spearman correlation value and a bottom 5 reconstruction loss.

**Figure S3.**
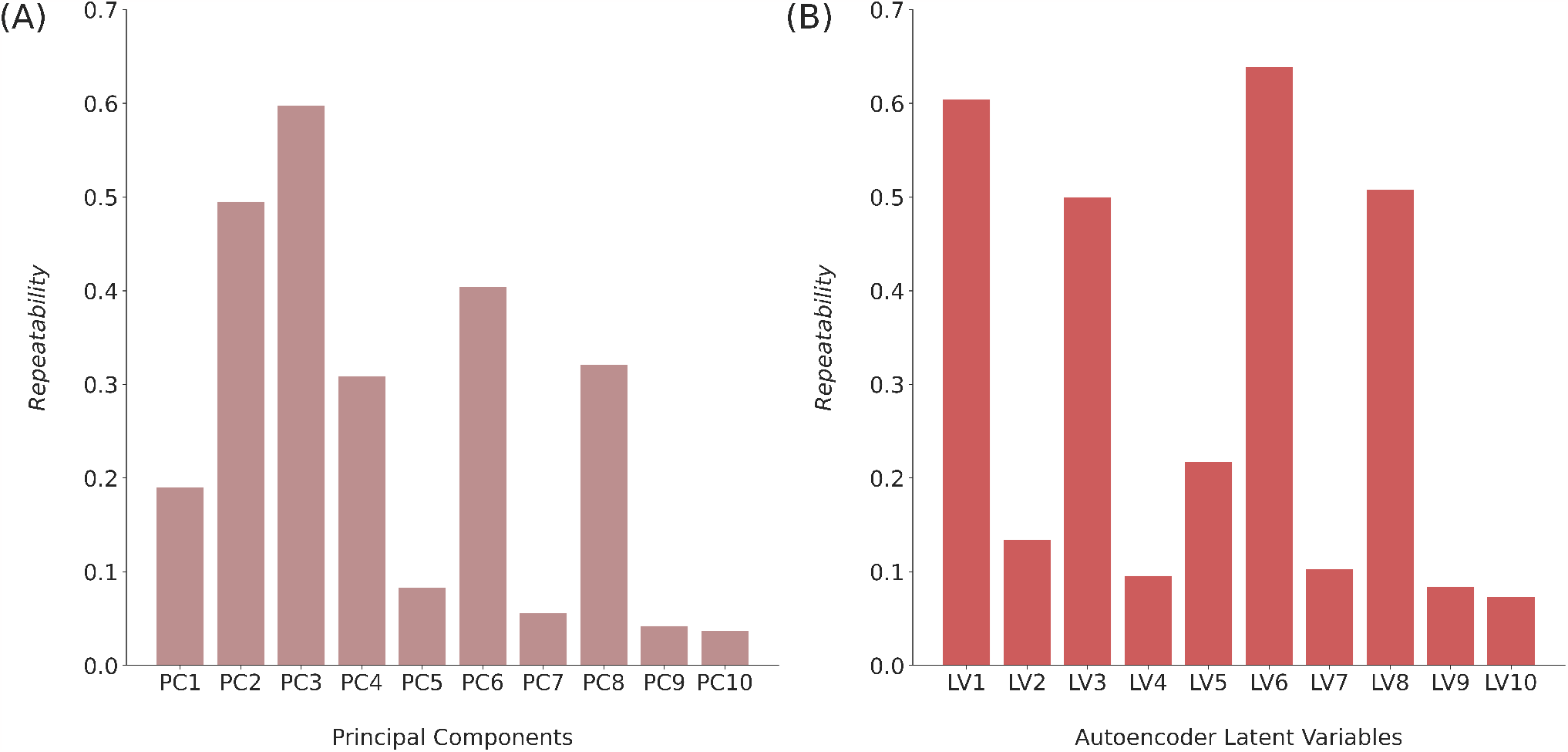
Comparison of repeatability values for reduced dimensions via PCA and an autoencoder neural network. (A). Repeatability ie. the proportion of total variance explained by genetics, represented on the Y-axis for each principal component (PC) on the X-axis. (B). Repeatability represented on the Y-axis for each autoencoder latent variable (LV) on the X-axis.

**Figure S4.**
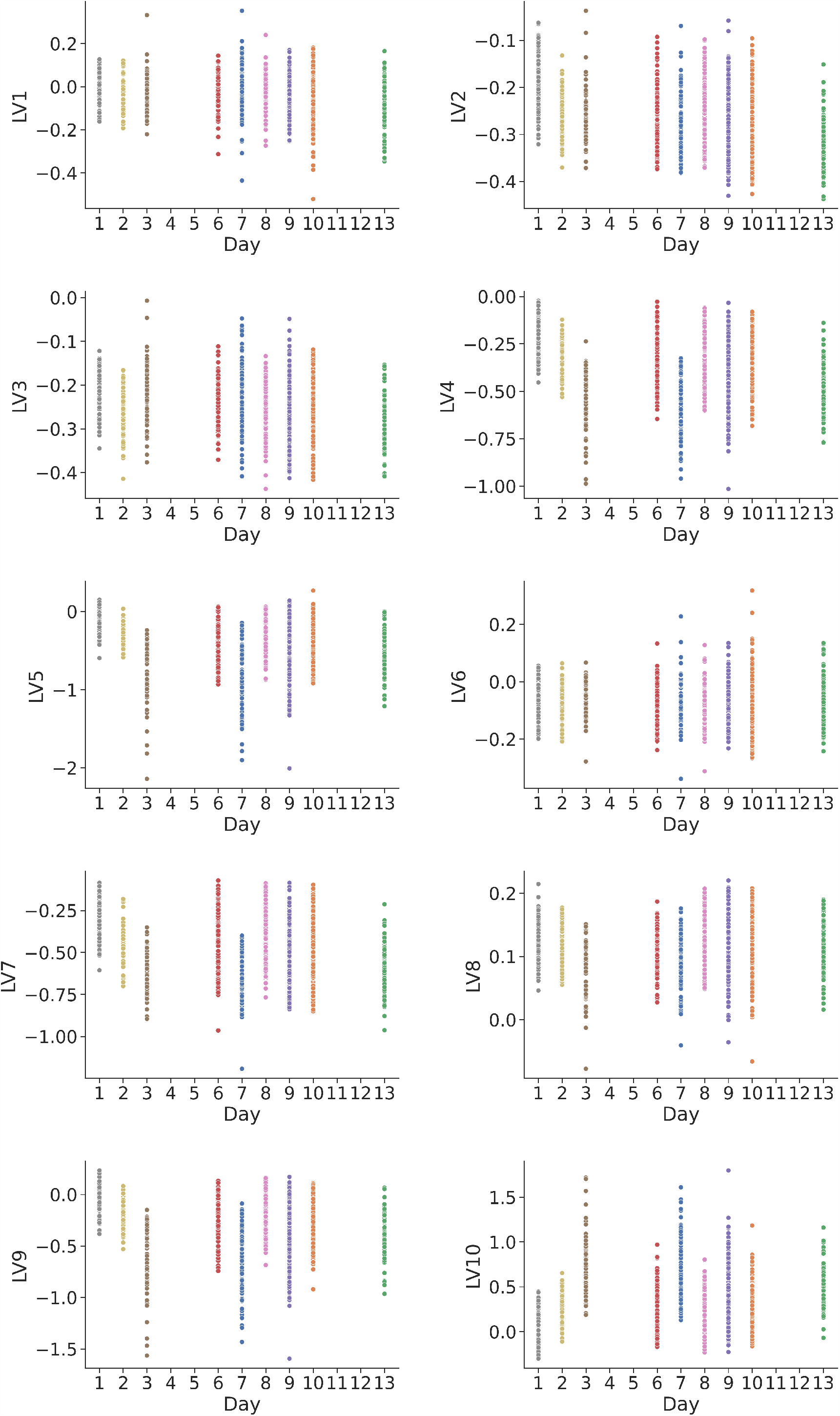
Association of auto-encoder derived latent variables of hyperspectral leaf reflectance values with day data of data collection for each individual plot. Y-axis of each subplot indicates the value for 1 of 10 latent variables. X-axis of each subplot indicates the day of data collection.

**Figure S5.**
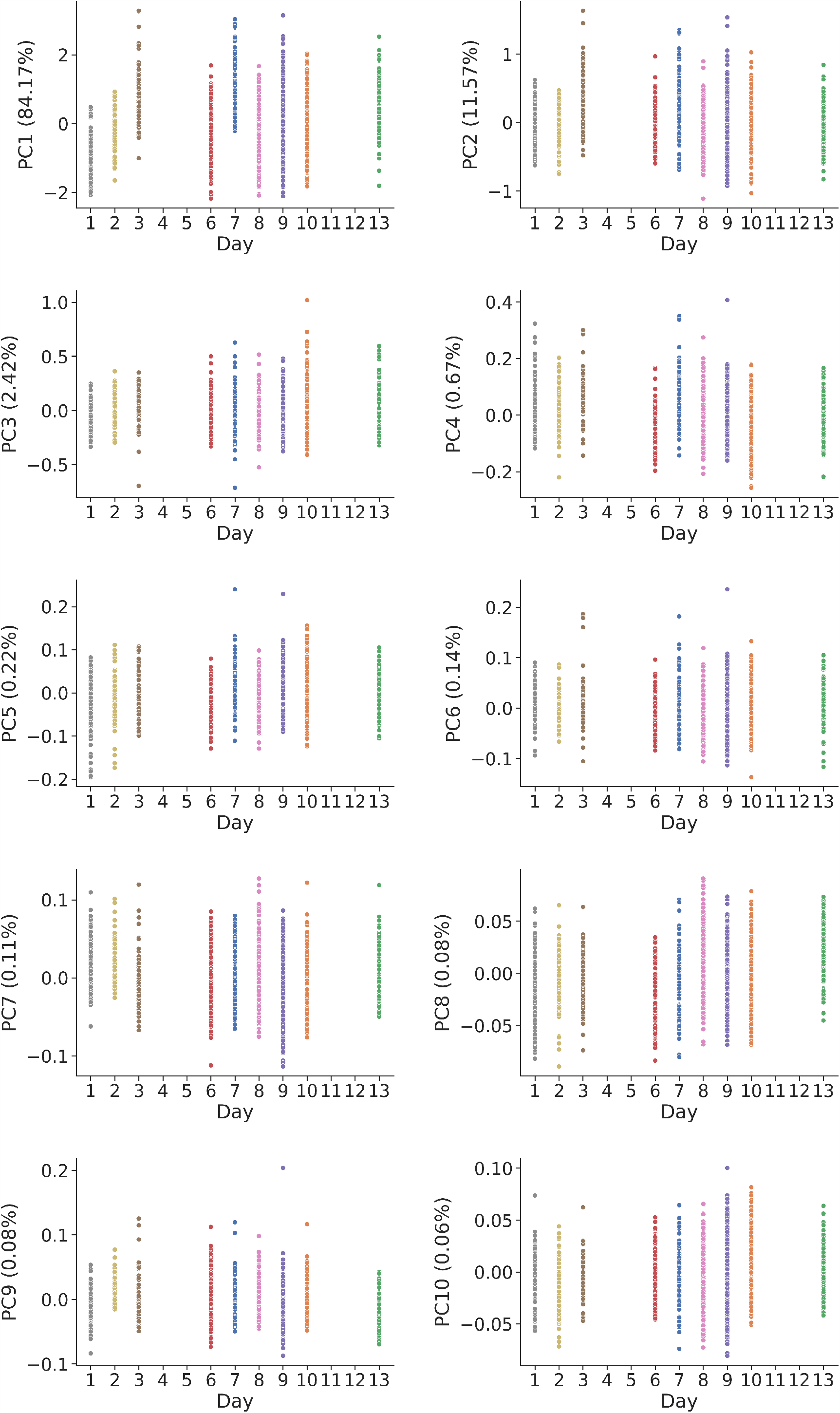
Association of principal components of hyperspectral leaf reflectance values with day data of data collection for each individual plot. Y-axis of each subplot indicates the value for 1 of 10 principal component and the proportion of variance explained by that component. X-axis of each subplot indicates the day of data collection.

**Figure S6.**
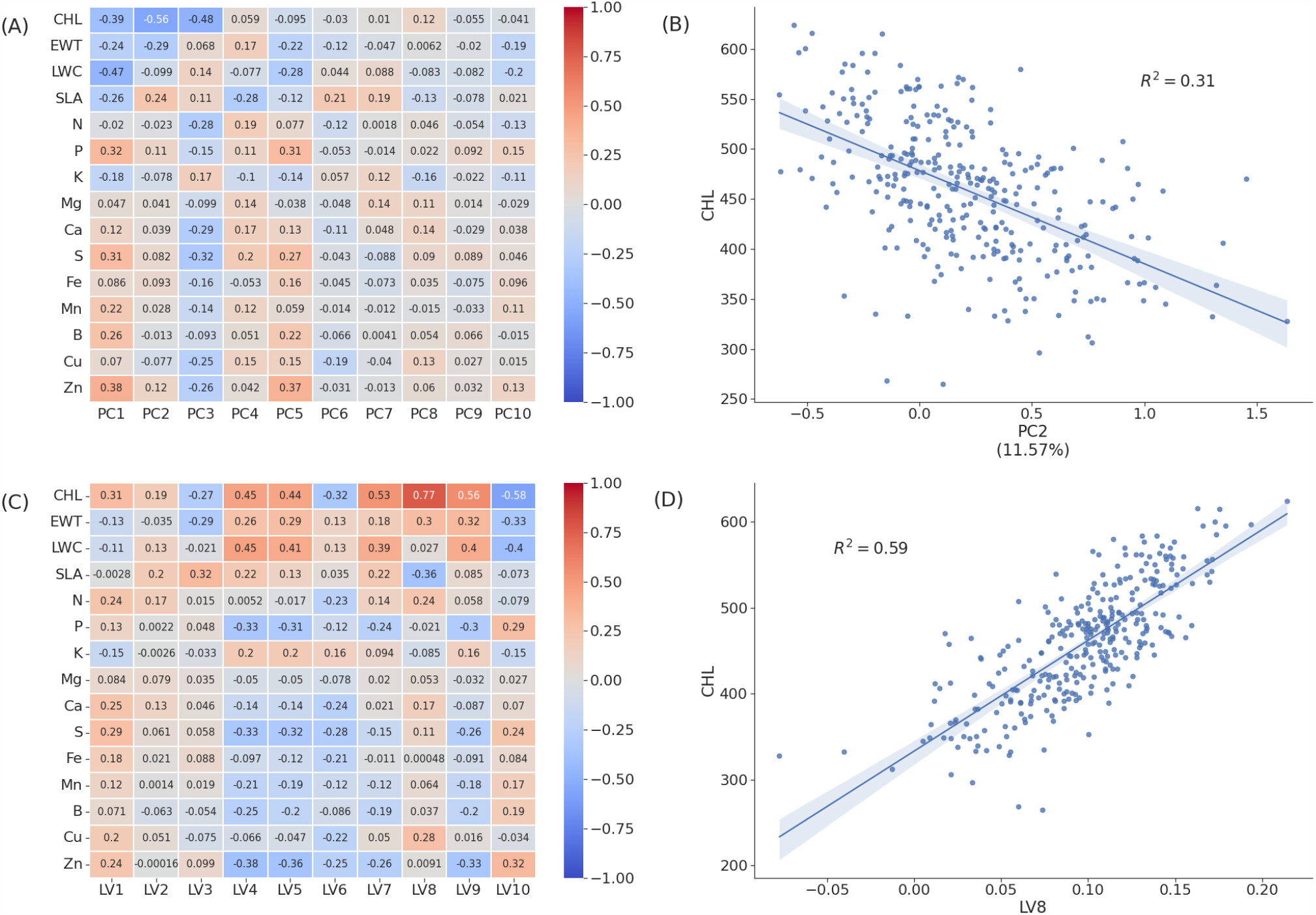
Associations between molecular traits and variables derived from autoencoder neural network (latent variables) and principal component analysis (PCs). (A). Spearman correlation coefficients between individual principal components and ground truth measurements for 15 traits scored for 243-318 plots as part of this study. (B). Correlation between chlorophyll content and principal component 2 derived from the autoencoder. (C). Spearman correlation coefficients between latent variables and molecular traits estimated in this study. (D). Correlation between chlorophyll content and latent variable 8.

**Figure S7.**
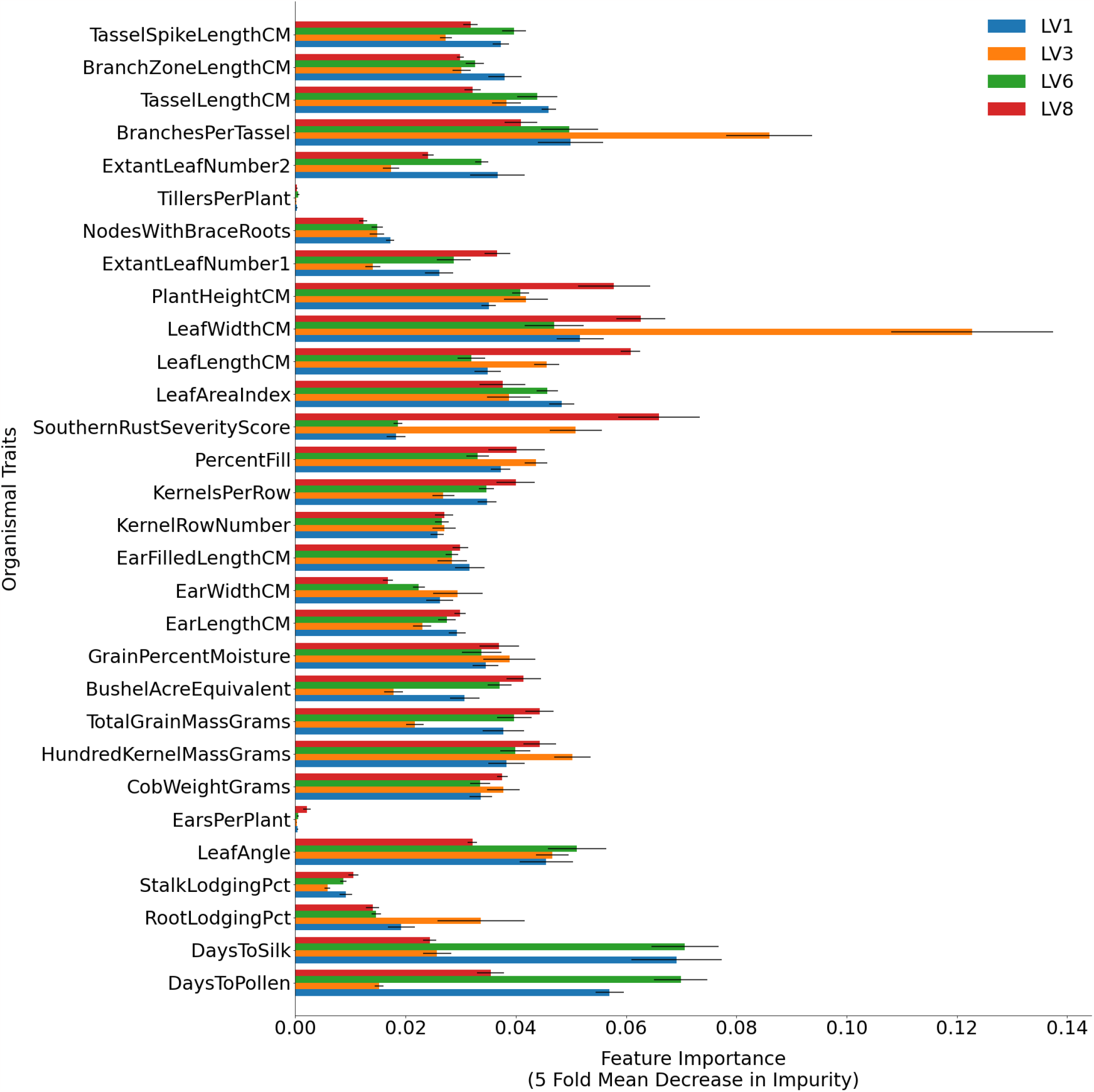
The relative importance of 30 hand measured traits in predicting latent variable 1, 3, 6, and 8 of the hyperspectral leaf reflectance values. This importance is derived from the mean decrease in impurity of each node in the decision trees within a trained random forest model, attributed to each feature (trait). X axis indicates the five fold mean decrease in impurity calculated for each trait. Error bars represent standard error across the five folds for each trait. Y-axis indicates the 30 hand measured traits used to predict the latent variable 3.

**Figure S8.**
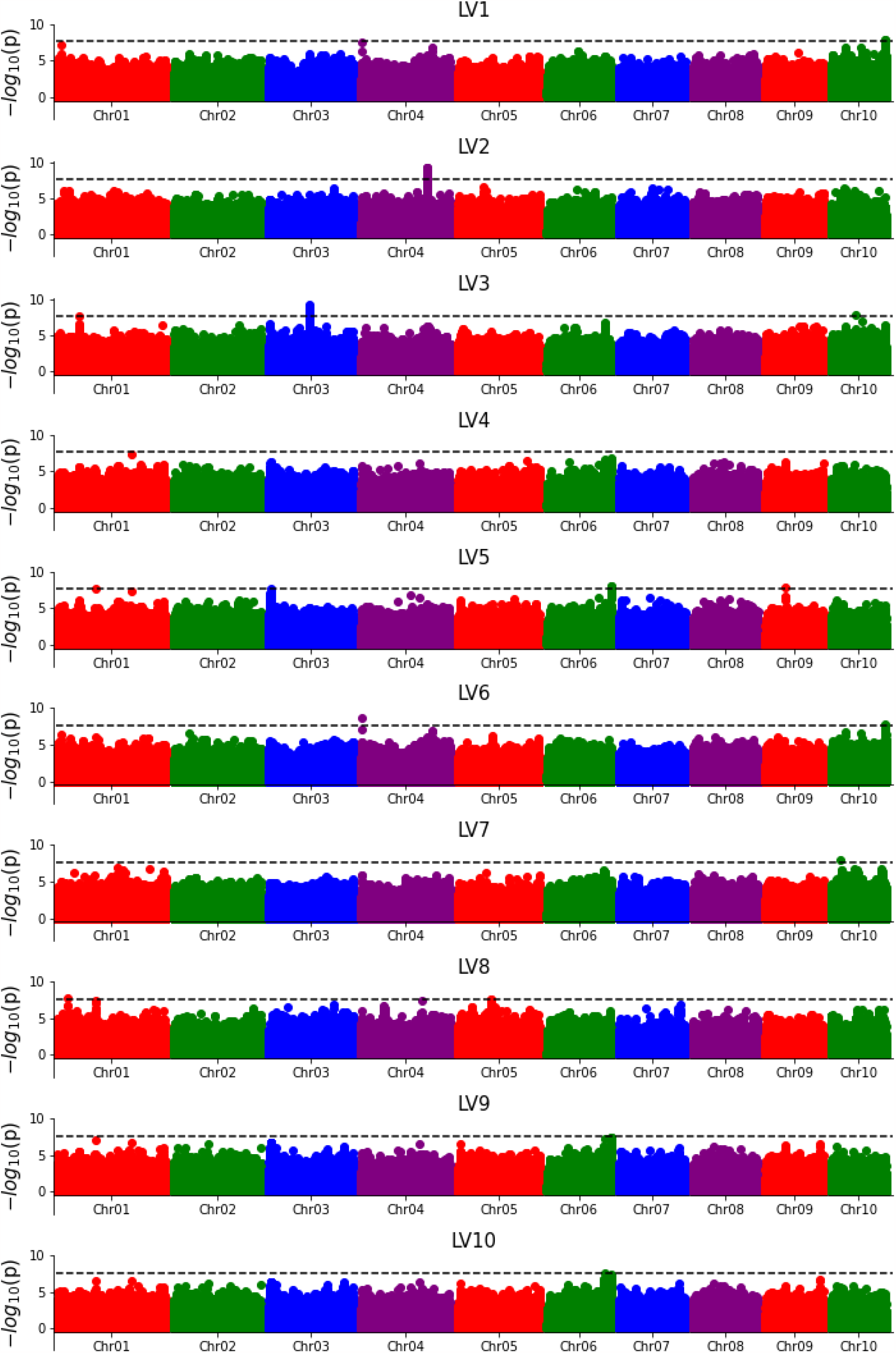
Genetic markers significantly associated with latent variables. Each point indicates the statistical significance of a marker (y axis) and it’s exact position on the genome (x axis). The dashed black lines indicate the statistical threshold cut off –2.20 x 10-8 – which was derived from an alpha value of 0.05 with a Bonferonni adjustment for the 2,269,711 effective number of markers.

**Figure S9.**
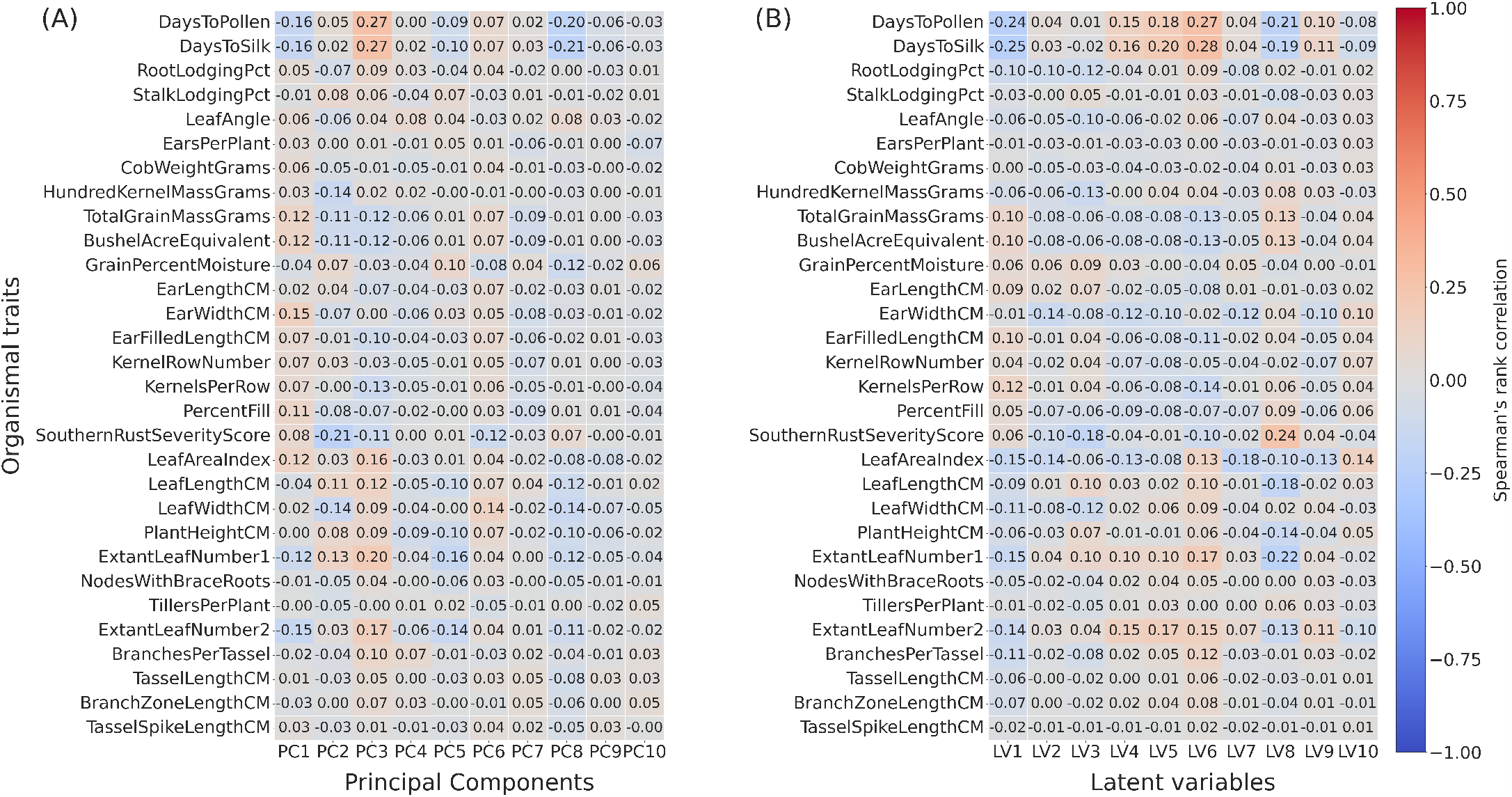
Associations between organismal (hand measured traits) and reduced dimensions via principal component analysis and auto-encoder neural network. (A) Spearman correlations between organismal traits (Y-axis) and principal components (PCs) (X-axis) of the hyperspectral leaf reflectance data. (B) Spearman correlations between organismal traits (Y-axis) and autoencoder derived latent variables (X-axis) of hyperspectral leaf reflectance data.

